# Stromal Reactivity Differentially Drives Tumor Cell Evolution and Prostate Cancer Progression

**DOI:** 10.1101/159616

**Authors:** Ziv Frankenstein, David Basanta, Omar E. Franco, Yan Gao, Rodrigo A. Javier, Douglas W. Strand, MinJae Lee, Simon W. Hayward, Gustavo Ayala, Alexander R.A. Anderson

## Abstract

We implemented a hybrid multiscale model of carcinogenesis that merges data from biology and pathology on the microenvironmental regulation of prostate cancer (PCa) cell behavior. It recapitulates the biology of stromal influence in prostate cancer progression. Our data indicate that the interactions between the tumor cells and reactive stroma shape the evolutionary dynamics of PCa cells and explain overall tumor aggressiveness. We show that the degree of stromal reactivity, when coupled with the current clinical biomarkers, significantly improves PCa prognostication, both for death and recurrence, that may alter treatment decisions. We also show that stromal reactivity correlates directly with tumor growth but inversely modulates tumor evolution. This suggests that the aggressive stromal independent PCa may be an inevitable evolutionary result of poor stromal reactivity. It also suggests that purely tumor centric metrics of aggressiveness may be misleading in terms on clinical outcome.

## Introduction

Stromal-epithelial interactions are well-established mediators of development in most organs, including common cancer sites such as breast, gastro-intestinal tract and prostate (Parmar and Cunha, 2004; Simon-Assmann et al., 2010; Strand et al., 2010). These interactions continue in adulthood where they maintain tissue differentiation and regulate growth. In cancer, alterations in the relationship between and within the stromal and epithelial tissues contribute to tumor growth and progression (Ayala et al., 2003; Bremnes et al., 2011; Franco et al., 2011; Kiskowski et al., 2011; Olumi et al., 1999; Orimo et al., 2005; Sugimoto et al., 2006; Tuxhorn et al., 2001; Tuxhorn et al., 2002b). The similarity between reactive stroma surrounding tumors and that surrounding wounds was observed more than 30 years ago (Dvorak, 1986). This includes activation of fibroblasts, recruitment of inflammatory cells, remodeling of the ECM and secretion of growth factors and cytokines. Experimental models have shown that carcinoma associated fibroblasts (CAF) are able to promote tumorigenesis in genetically-initiated benign epithelial cells. We have demonstrated the effects of stroma on epithelium, and also interactions between sub-populations of fibroblasts influencing stromal to epithelial signaling (Franco et al., 2011; Kiskowski et al., 2011; Olumi et al., 1999).

A number of studies have found that tumor stroma is genetically intact and stable despite being morphologically abnormal, presumably in response to altered local paracrine signals (Bianchi-Frias et al., 2016; Qiu et al., 2008; Weinberg, 2008). This suggests that stromal cells may be more amenable to therapeutic intervention than the genetically unstable tumor epithelium. It is also possible that adjusting or normalizing the interactions between stromal and epithelial cells may represent a mechanism for modifying tumor aggressiveness. A well-defined model of these interactions would facilitate a more focused search for new medical therapeutic approaches.

Many chemokines and cytokines play a documented role in tumor-promoting paracrine interactions (Franco et al., 2011; Grivennikov et al., 2010; Kiskowski et al., 2011; Raman et al., 2007). However, the mechanisms of the signaling milieu surrounding tumors are complex and the fine details are not well understood. Dissecting the dialogue between the tumor and the stroma is a major experimental undertaking that would be considerably simplified by the development of integrated mathematical/experimental approaches that focus on cellular consequences rather than individual genes. The cellular phenotype is a product of both genetic and non-genetic determinants and is always defined in relation to a specific context. Cellular selection occurs at the phenotypic level, and this is the scale which naturally integrates both intrinsic and extrinsic signals to produce a functional response.

We have focused on prostate cancer as a model of carcinogenesis. Prostate cancer (PCa) is a significant health care problem due to its high incidence and mortality (Penson et al., 2003; Potosky et al., 2004). The disease has a wide and varied clinical spectrum. Most PCas that are diagnosed early have an indolent course. In contrast, a sub-group of PCas at diagnosis are so aggressive that surgery or radiation cannot control them (Amling et al., 2000; Han et al., 2001; Hull et al., 2002; Moul, 2006; Pound et al., 1999; Roehl et al., 2004). Current prognostic standard of care approaches are limited in their ability to predict these individual patterns of progression, resulting in a high level of overtreatment (Harrington et al., 2016). Cancer-based biomarkers continue to be imperfect at identifying patients who present with localized disease but are likely to fail standard treatment and progress to fatal disease.

Reactive stroma initiates early in the pre-malignant prostatic intraepithelial neoplasia (PIN) and promotes prostate tumor growth (Tuxhorn et al., 2001; Tuxhorn et al., 2002a; Tuxhorn et al., 2002b). Reactive stroma grading (RSG) is an independent predictive factor for PCa biochemical recurrence and PCa specific death (Saeter et al., 2015, 2016) and can add significant predictive value to Gleason grading (McKenney et al., 2016). We have defined stromogenic carcinoma as lesions containing reactive stroma in >50% of the tumor (Ayala et al., 2003; Yanagisawa et al., 2008). The presence of stromogenic carcinoma can predict biochemical recurrence in needle biopsies (Yanagisawa et al., 2008) and the percentage of stromogenic carcinoma in the entire tumor in radical prostatectomy specimens can predict biochemical recurrence and PCa specific death (Ayala et al., 2011). Reactive stroma has also been correlated with tumor progression in many other cancers, such as lung, breast, and skin (San Martin et al., 2014).

There is a small but growing literature on mathematical models of tumor-stroma interactions, driven by our group (Basanta et al., 2012; Basanta et al., 2009; Flach et al., 2011; Kim et al., 2013) and also by others (Kim and Othmer, 2013; Martin et al., 2010). However, none of these considers how stromal activation may differentially alter tumor evolution. Pertinent to our current work, is a previous model which employed a hybrid cellular automaton (HCA) approach to characterize the glandular architecture of prostate tissue and its homeostasis through a layered epithelial homeostasis via growth factor signaling regulated by surrounding stroma (Basanta et al., 2009). Thus far none of these previous works have consider the impact of stromal reactivity on tumor evolution.

We initiated the current study using pathologic features of human prostate cancer, and have integrated mathematical and biological modeling to focus on phenotypes that elucidate disease initiation and local invasion. We developed a multiscale mathematical model that has suggested novel hypotheses regarding prostate cancer progression. Specifically, we investigate how the interplay between stromal components and a heterogeneous tumor epithelium modulate tumor evolution, growth and invasiveness. The predictions generated by this model were tested in biological models and cross-validated in large cohort of human samples. We discovered that PCa stroma exerts selection pressure that drives cancer, heterogeneity, growth and regulates the evolution to a lethal phenotype. Perhaps even more profound, the overall behavior of the tumor in patients cannot be accurately predicted by either the cancer cell or the stromal response alone. In fact, the model predicts that the degree of stromal reactivity, when integrated with the current clinical methodology (Gleason grading with clinico-pathologic parameters), significantly improves PCa prognosis predictions and may be used to modify treatment decisions. This is in concordance with a recent clinical study (McKenney et al., 2016). Counter-intuitively our mathematical model also predicts that the degree of PCa stromal reactivity inversely correlates with evolution of the cancer cell population to more aggressive phenotypes. This suggests that the aggressive stromal-independent PCa may be an evolutionary result of poor stromal reactivity.

## Results

### Multiscale prostate peripheral zone model characterizes the tumor-microenvironment dialog

In order to understand the role of the tumor microenvironment in prostate cancer progression, we developed a model using a Hybrid Cellular Automata (HCA) paradigm (Anderson, 2005; Quaranta et al., 2005; Rejniak and Anderson, 2011) that recapitulates this interplay. We designed the model to explicitly capture cellular phenotypic heterogeneity within the context of a dynamic spatio-temporal microenvironment. Each cell is considered as an individual and its behavior emerges as a function of its phenotype under the influence of its local microenvironment. We consider 6 mathematically abstracted cell types: normal basal and luminal epithelial cells; tumor epithelium; native stroma (e.g. muscle, fibroblasts); reactive stroma (i.e. CAF/myofibroblasts) induced from normal stroma; and motile stroma (representing a generic cell with inflammatory properties). The physical microenvironment is described by a system of continuum deterministic equations (see methods), which includes growth factors (GF; affecting cell proliferation and function), basement membrane (BM; representing a mechanical barrier to the glands), extracellular matrix (ECM; structural support to intercellular communication and growth), matrix metalloproteinases (MMP; degradation of the ECM), and empty space (assumed to be occupied by interstitial fluid) (Figure 1A). These equations are coupled in space and time on a two-dimensional lattice, which is based on a histological slide of the human prostate peripheral zone (Figure 1B-D).

**Figure 1.**
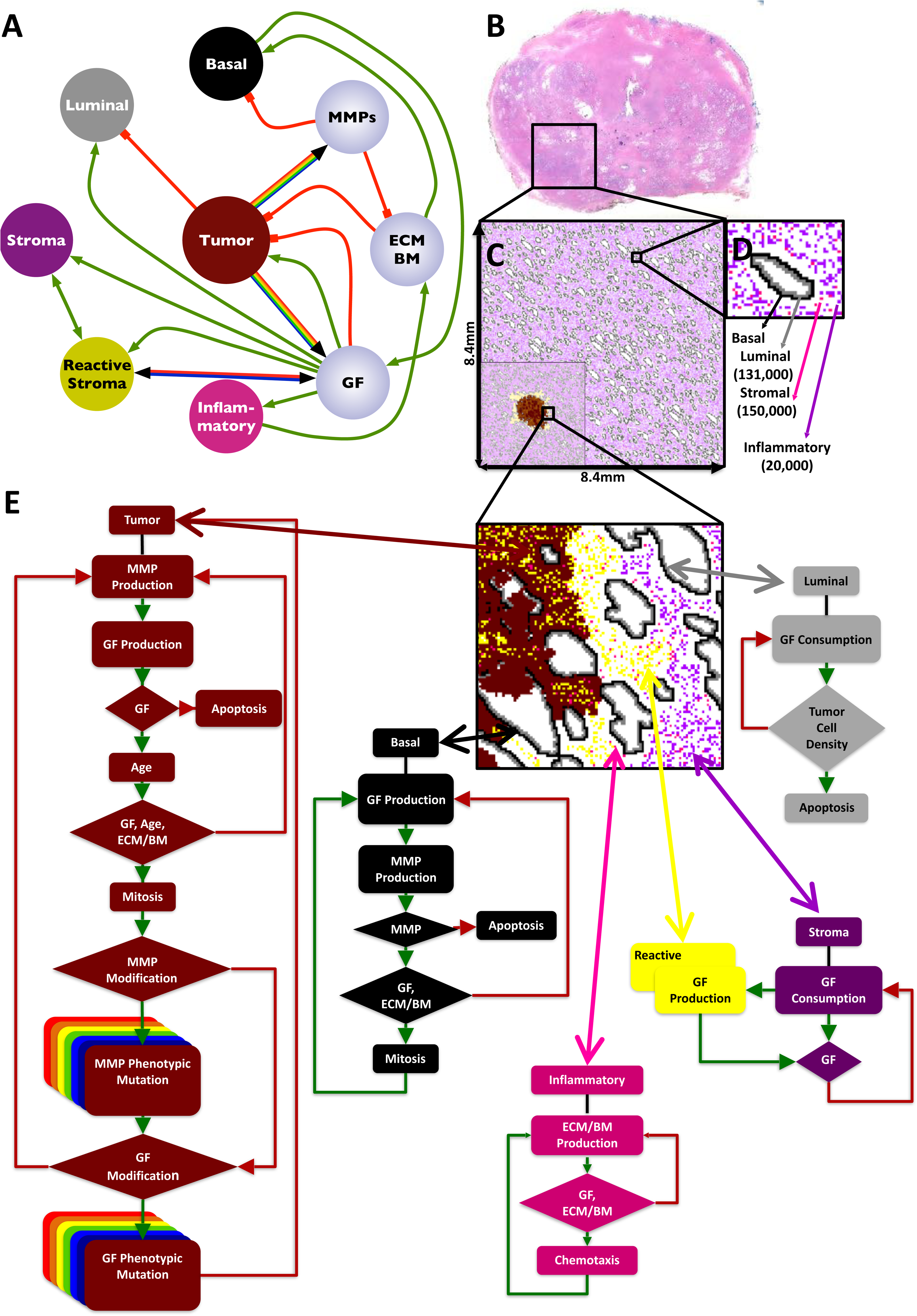
In silico multiscale model of the prostate peripheral zone. Interaction network of key model variables (A). Interactions between cells (colored nodes) and microenvironmental variables (lilac nodes) are represented as either green (positive) or red (negative) connections. Multicolor connectivity represents the spectrum of possible tumor phenotypes with different levels of GF and MMP production. Bicolor connectivity represents two different degrees of Stromal Reactivity. In silico reconstruction of the normal prostate peripheral zone tissue (B-D). Histopathological slide of the whole normal prostate, highlighting the peripheral zone, filled with epithelial acini surrounded by stroma (magenta) (B). In silico representation of the complete peripheral zone, including ductal structures and cellular densities that mimic normal anatomy (C). This constitutes the domain where all simulations were performed. Inset figure on the bottom left is an example of a sample simulation Representation of a single reconstructed duct and the surrounding stroma as well as the total number of cell types. (D). Cell decision flowcharts for each cell type in the model (E). The phenotypic behavior of an individual cell is based on the interaction between the cell and the local microenvironment.

We used a histological slide from a normal human prostate to generate a baseline model of the peripheral zone, within which a tumor is initiated. Image segmentation was used to retrieve the basic anatomical glandular structures and cellular densities (Figure 1B). To reconstruct the tissue domain, histologic information was discretized on a two dimensional lattice (Figures 1C and 1D). PCa pathogenesis was simulated by seeding a single abnormal luminal cell inside a duct near the center of the lattice. Typically, the tumor cell populates the duct through division and then breaches the basement membrane. The tumor mutates and appropriate phenotypes invade throughout the peripheral zone, eventually reaching the edge of the lattice. In our model, as shown in supplementary figure S1, cells are capable of migration to orthogonal positions. Cells are also capable of division, apoptosis as well as production and consumption of extracellular proteins and growth factors, as dictated to by the cell lifecycle flowcharts (Figure 1E).

To study the impact of environmental selection on prostate cancer phenotypic heterogeneity, eight initial tumor phenotypes with different levels of GF and MMP production were chosen. These are key biological drivers of prostate cancer growth and invasion. Tumor cells in the model mutate randomly (through unbiased drift at division) from their parental phenotype by altering their GF and MMP production rates. To explore the role of stroma, two different stromal reactivity phenotypes were modeled: high stromal reactivity (high SR; stromal cells that upon activation produce high amounts of GF) and low SR (stromal cells that upon activation produce low amounts of GF). This leads to 16 different combinations of tumor-stromal growth conditions. We performed a total of 4,800 simulations consisting of 300 repeats for the 16 different tumor-stromal combinations.

### Cancer-stroma phenotypic dialogue regulates tumor growth and invasion in a non-linear manner

*In silico*, the level of stromal activation, defined as the proportion of stromal cells activated per year, is significantly correlated with the level of tumor growth (tumor epithelium and reactive stroma). Representations of tumors growing in low and high SR are presented (Figure 2A and B). Tumors, initiated using a single cancer cell characterized by low GF and MMP production, grow faster in microenvironments with higher SR. This suggests that the degree of SR may be at least as important a driver of differential tumor growth and invasion as the phenotype of the initially seeded tumor cell (Figures 2C and 2H). More specifically, tumors grow faster (as measured by time to maximal size, i.e. time to edge of domain) not only when initiated with an aggressive epithelial phenotype (i.e. high levels of GF and MMP production) but also when seeded with non-aggressive phenotypes, provided the tumor microenvironment contains high SR (Figure 2D). These data are consistent with research from our labs and others that have emphasized the role of reactive stroma facilitating tumor growth (Fluge et al., 2009).

**Figure 2.**
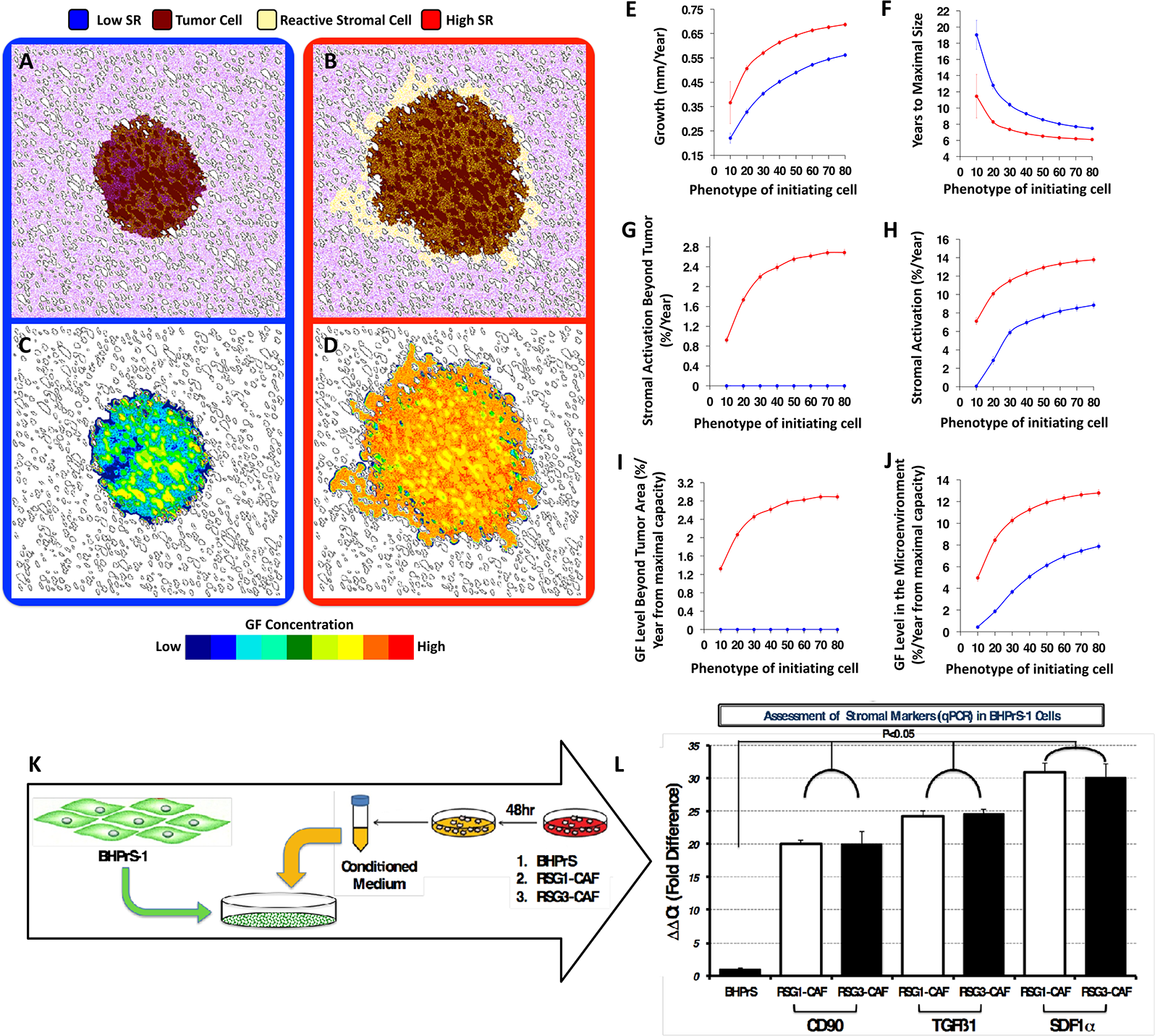
Change in SR phenotypes, tumor growth and invasiveness. 6 years of simulated tumor growth, initialized with a tumor cell producing low levels of GFs (20% of the maximal simulated cell production capacity); under two different microenvironmental conditions: (A, C) low SR (blue frames and lines), or (B, D) high SR (red frames and lines). Different tumor metric distributions over 8 initiating phenotypes (ranging from 10-80% of the maximal simulated cell GF production capacity) in high (red) and low (blue) SR environments averaged over 100 simulations per phenotype: Average growth of the tumor in mm per year (E); Time to achieve maximal size (reach the edge of the tissue domain) (F). Percentage of stromal activation within (G) and beyond (H) the tumor varies with the phenotype of the tumor initiating cells and with the stromal reactivity. The concentration of GF found in the microenvironment beyond (I) and within (J) the tumor parallel the stromal activation. To assess the ability of reactive stroma to activate benign stroma cells the human prostate stromal cell line BHPrS1 was cultured in the presence of conditioned medium (CM) from either BHPrS1, RSG1-CAF or RSG3-CAF for 4 weeks (K). At the end of the experiment expression of CD90, TGFß1 and SDF1 (all putative activated stromal markers) was determined by qPCR. Conditioned media from RSG1- and RSG3-CAF elicited a similar and significant increase in the levels of these mRNAs compared to medium conditioned by the functionally normal BHPrS1 cell line (L).

This model also makes the intriguing prediction that stromal cells with high reactivity can be activated not only within or adjacent to the tumor, but also at some distance beyond the tumor margins (Figure 2B and E). This phenomenon arises from the GF production initiated by the tumor cells, which activates local stromal cells. Once triggered, the paracrine production of GF by high-SR will form an autonomous activation cascade that extends beyond the edge of the tumor (Figure 2H and I). In the case of low SR, reactive stromal cells are only found within the tumor, as they are much more dependent on the GF produced by tumor cells directly (Figure 2F and G). To test this idea we exposed a benign human prostate fibroblast cell line (BHPrS1) to conditioned medium from either RSG1 or RSG3-derived carcinoma associated fibroblasts (Figure 2K). We tested for expression of three key markers found associated with stromal activation (CD90, TGF-ß1 and SDF1α) and compared these expression in BHPrS1 cells growing autonomously conditioned medium. Both RSG1-CAF and RSG3-CAF conditioned medium were able to cause upregulation of stromal activation markers in this system, no such upregulation was seen with conditioned medium from benign stromal cells.

In summary, differential tumor growth and invasion is correlated with the levels of GF production by the initiating tumor cell and the degree of SR. However, this is a non-linear relationship (similar to Michaelis-Menten kinetics). The results from the *in silico* model also predict that the degree of SR is reflected by the proportion of activated stromal cells. Taken together, this suggests that the phenotypes of the tumor cells and the degree of SR could, in combination, act as a more accurate prognostic marker.

### **Human-derived stroma drives cancer growth *in vivo* in a stromal grade-specific manner.**

To examine the prediction that the nature of reactive stroma plays a role in tumor promotion we isolated and validated CAF from human prostate cancer samples as previously described (Olumi et al., 1999). Sections of the source tissues were scored for reactive stromal grade according to standard guidelines (Figure 3A), (Ayala 2003). Stromal grading was done without regard to the Gleason score in these samples (Figure 3B). Analysis of the Ki67 index revealed increased proliferation in cancer cells with RSG3 compared to RSG1 (Figure 3B) in this model.

**Figure 3.**
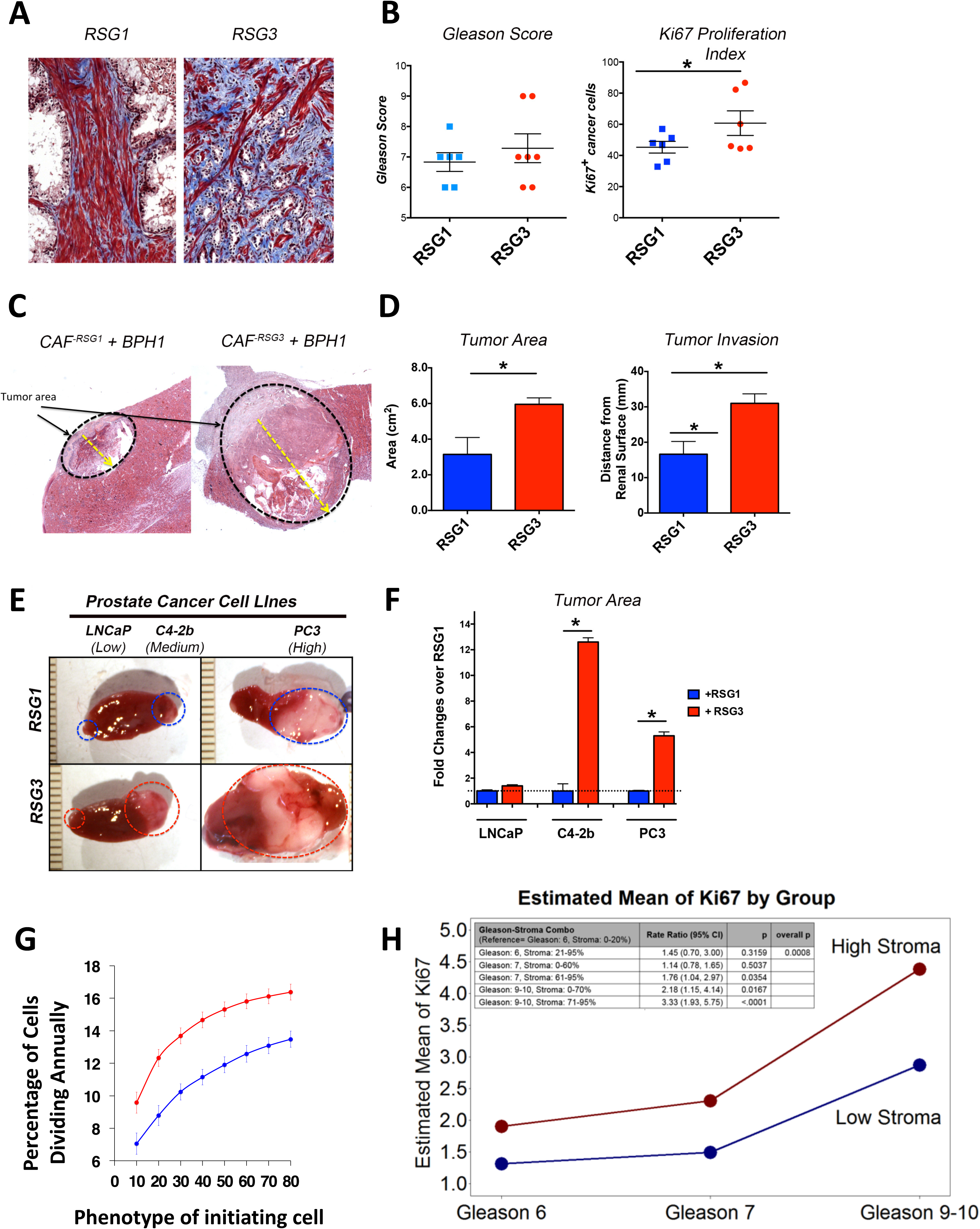
In-vivo stromogenic grade and auto reactivity is linked to tumor growth and invasion but not to Gleason grade. Representative images of stromogenic response in prostate cancer from two different patients showing RSG1 (left) and RSG3 (right). Note the intense and high percentage of reactive stroma (blue) depicting increased collagen deposition in the RSG3 sample (A). A total of 23 patients were categorized according to their Gleason Score (left). No correlation was found between Gleason score and RSG. However, cancer cells surrounded by RSG3 stroma had increased proliferation compared to RSG1 (B). Response of an initiated reporter epithelial cell line (BPH-1) to CAF is a function of the RSG status of the tumor source of the CAF. Low magnification of CAF combined with BPH1 cells in vivo. Both CAF-RSG1 and CAF-RSG3 promoted malignant transformation (C). Quantitation of tumor area and invasion revealed increased growth and aggressiveness in CAF-RSG3 compared to CAF-RSG1 (D). Response of epithelial cells to CAF is a product of both the epithelial and stromal components of the tumor. Gross picture of PCa cell lines LNCaP, C4-2b and PC3 cells tissue recombinants with RSG1 and RSF3 CAF (E). Fold change analysis of PCa cell lines combined with RSG CAFs TR shows significant increased growth in the presence of RSG3 compared to RSG1 in C4-2b and PC3 but not with LnCaP cells (F). Tumor cell division rate calculated from our in silico model simulations over 8 different initiating phenotypes in high (red) and low (blue) SR environments (averaged over 100 simulations) (G). The proliferation rate of cancer cells, as measured by Ki67, is significantly higher in stromogenic than non-stromagenic cancers, in all Gleason categories (H).

Recombinants were generated using an initiated but non-tumorigenic prostate epithelial cell line, BPH1 previously validated as a reporter of the tumor-inducing effects of CAF (Ao et al., 2007; Franco et al., 2011; Olumi et al., 1999). CAF alone do not form tumors. In recombinants CAF from each RSG group induced tumors *in vivo* (Figure 3C). Quantitation of tumor area and invasion revealed that recombinants made using CAF-derived from RSG3 patient tumors were significantly larger and more invasive compared to RSG1-derived cells (Figure 3D). This demonstrates that stromal characteristics, and in particular the extensive stromal response seen in RSG3 can be a powerful driver of tumor growth and local invasion.

To further investigate whether the nature of the epithelium modifies the effects of CAF we utilized three human prostate cancer lines as reporters. Using the same tissue recombination model applied to BPH1 cells we showed that responses were indeed epithelial cell type specific. Tumor growth was a function of both the aggressiveness of the epithelial cells and the stromogenic status (RSG1 vs. RSG3) of the source of the CAF. Consistent with *in silico* model predictions, RSG3-derived CAF induced significantly larger tumors than RSG1 CAF in C4-2B (>12 fold) and PC3 (>6 fold) cells. These lines are both tumorigenic and metastatic when grafted alone, as compared to the LNCaP line from which the C4-2B is derived, which result in small tumors when grafted alone. While the recombinants using RSG3 CAF and LNCaP were slightly larger than their RSG1-derived CAF counterparts this was a small and statistically insignificant change (Figures 3E and 3F). Thus the response of the epithelial cells seems to be broadly correlated with their degree of aggression, where BPH1 cells alone are non-invasive, as are LNCaP, while both PC3 and C4-2B cells are aggressive and invasive, characteristics that are enhanced by the stromal grade of the source of associated fibroblasts.

Based on the *in vitro* Ki67 analysis (Figure 3B) we calculated the percentage of tumor cells that were undergoing division per year in our *in silico* model under the 2 differential stromal conditions and the 8 initiating cancer cell phenotypes. Figure 3G shows a consistent increase in the fraction of proliferating tumor cells when using high SR, compared to low SR. This prediction was further tested in a large cohort of human patient samples. The results show that tumors with RSG3 have a consistently higher proliferation index when analyzed for every Gleason category (Gleasons 6, 7, 8-10) (Figure 3H). These data support the in vivo data and validate our approach.

### A novel Integrated Cancer Biomarker (ICB): Combination of Reactive Stroma Grading and Gleason

Based on the results obtained *in silico* and *in vivo*, we created a clinical combination biomarker with data from the epithelium and stroma. As a stromal marker we used the percentage of RSG3 within radical prostatectomy PCa, as previously described (Ayala et al., 2011; Li et al., 2009; Li et al., 2004; Maru et al., 2001). We chose Gleason score as an epithelial cancer marker. This new “Integrated Cancer Biomarker” (ICB), was tested against biochemical recurrence and PCa specific death. Optimal cut-off values for percentage of RGS3 were obtained by using minimum p value approach for every Gleason category (Figures 4A-F). Gleason 6 patients were separated with a cutoff of 21% reactive stroma and above (RSG-GS6) (histology in figure 4A and 4B); Gleason 7, 61% and above (RSG-GS7) (Figure 4C and 4 D) and Gleason 8-10 with 71% and above (RSG-GS8-10; Figure 4E and 4F).

**Figure 4.**
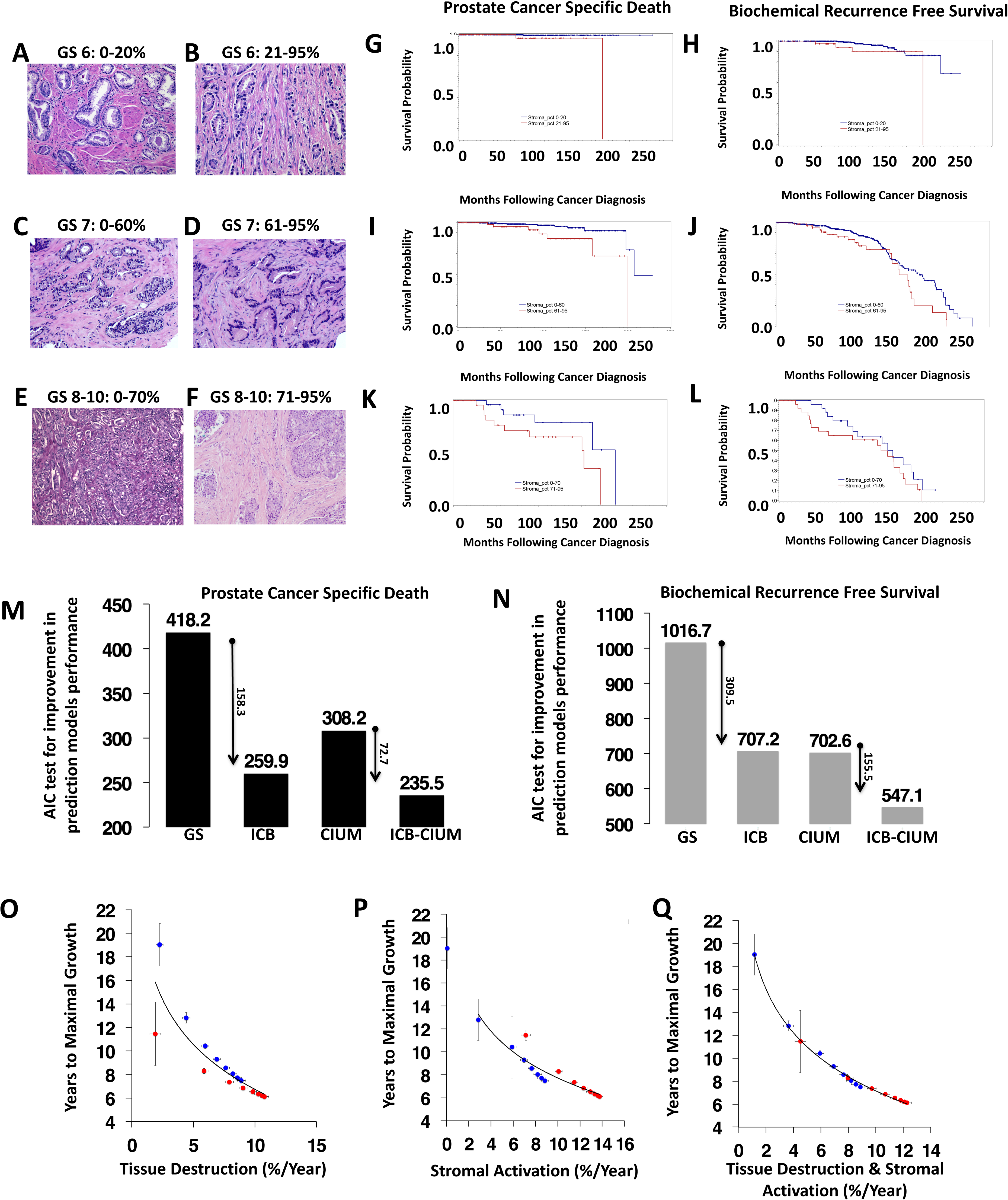
A novel Integrated Cancer Biomarker (ICB) stratifies all Gleason grades in a cohort of 1,291 prostate cancer patients with over 20 years of follow-up. Histology of a Gleason 6 cancer without stromal response (A) compared to a Gleason 6 with exuberant stromogenic response (RSG3) (B). Patients with more than 20 % of the tumor having a RSG3 pattern are associated with increased PCa specific death (G) and biochemical recurrence (H). Histologic representations for Gleason 7 cancers without reactive stroma (C) and with RSG3 in (D). In Gleason 7 patients the cutoff for statistical significance is 60% of the tumor having a RSG3 pattern (I for PCa specific death and J for biochemical recurrence). High Gleason cancers (GL. 8-10) can grow in solid masses (E) or embedded in RSG3 (F). The cutoff in this category is higher, at 70% (K for PCa specific death and L for biochemical recurrence). Akaike information criterion (AIC) test comparing different predictive models of prostate cancer specific death and recurrence free survival: GS (Gleason Score), ICB (Integrated Cancer Biomarker), CIUM (Clinical In Use Methodology: GS, seminal vesicle invasion, extra capsular extension and PSA). The Aikake model for PCa specific death free survival (M) and biochemical recurrence free survival (N) show the much-improved performance of the Integrated Cancer Biomarker compared to standard of care. The preferred model is the one with the lowest AIC value. In Silico model predicted logarithmic correlation of time to maximal growth (in years) with different tissue metrics over 8 different initiating phenotypes in high (red) and low (blue) SR environments: (O) Normal tissue architecture destruction, (P) Stromal activation, and their combination (Q).

We used the Cox proportional hazard models to evaluate univariable and multivariable associations of time to death or recurrence with ICB. The hazard rates of each model were all compared to the reference group: patients with a Gleason score of 6 and 0-20% of reactive stroma grade RSG3 within the tumor (Table 1). Extra-capsular extension, seminal vesicle invasion, margins, lymph node status and preoperative PSA were adjusted in the final multivariable model. Those with GL6/21-95% of the tumor with RSG3 had a 2.4 fold increased risk for biochemical recurrence (BR) (Figure 4H) and 7.9 fold increased risk of PCa-specific death (PCaSD) (Figure 4G) than baseline. In the Gleason 7 category, those with 0-60% of the tumor with RSG3 had a 3.2 and 5.6 fold risk of BR (Figure 4J) and PCaSD (Figure 4I), while those with over 61% had 4.3 and 12.7 fold risk of BR (Figure 4G) and PCaSD respectively (Figure 4J). Even within Gleason 8- 10 categories, those with 0-70% of the tumor with RSG3 had a 3.2 and 18.6 fold risk of BR and PCaSD, while those with over 71% had 7.3 and 56 fold risk of BR and PCaSD respectively (Figures 4L and 4K). Note the difference that the increased RSG has on the hazard ratios in every category, these were highly significant (p < 0.0001) (Table 1).

**Table 1.** Multivariable association between ICB and risk of BR and PCa SD based on Cox PH model

| Variable | Adjusted Hazard Ratio<br>(95% Confidence Interval) | p-value |
| --- | --- | --- |
| **[Biochemical recurrence]** |  |  |
| **Integrated Cancer Biomarker (ref: GL 6 / RGS3 0-20)** |  | **<.0001** |
| GL 6 / RGS3 21-95 | **2.41 (0.67, 8.62)** | **0.1763** |
| GL 7/ RGS3 0-60 | **3.2 (1.70, 6.04)** | **0.0003** |
| GL 7 / RGS3 61-95 | **4.32 (2.05, 9.10)** | **0.0001** |
| GL 8-10 / RGS3 0-70 | **3.25 (1.42, 7.44)** | **0.0054** |
| GL 8-10 / RGS3 71-95 | **7.29 (3.37, 15.77)** | **<.0001** |
| Extra-capsular extension vs. no | 1.50 (0.95, 2.38) | 0.0834 |
| Seminal vesicle invasion vs. no | 1.35 (0.93, 1.96) | 0.1165 |
| Margins vs. no | 2.24 (1.55, 3.24) | <.0001 |
| Lymph node status vs. no | 2.82 (1.91, 4.16) | <.0001 |
| log(preoperative PSA) >1.9 vs. ≤1.9 | 1.80 (1.19, 2.72) | 0.0056 |
| **[PCa specific death]** |  |  |
| **Integrated Cancer Biomarker (ref: GL 6 / RGS3 0-20)** |  | **<.0001** |
| GL 6 / RGS3 21-95 | **7.89 (0.47, 131.5)** | **0.1499** |
| GL 7/ RGS3 0-60 | **5.64 (0.68, 46.58)** | **0.1083** |
| GL 7 / RGS3 61-95 | **12.69 (1.38, 117.0)** | **0.025** |
| GL 8-10 / RGS3 0-70 | **18.6 (1.85, 186.8)** | **0.013** |
| GL 8-10 / RGS3 71-95 | **55.82 (6.22, 501.0)** | **0.0003** |
| Extra-capsular extension vs. no | 1.94 (0.52, 7.33) | 0.3265 |
| Seminal vesicle invasion vs. no | 2.81 (1.26, 6.26) | 0.0118 |
| Margins vs. no | 1.26 (0.50, 3.14) | 0.6242 |
| Lymph node status vs. no | 1.84 (0.83, 4.05) | 0.1321 |
| log(preoperative PSA) >1.9 vs. ≤1.9 | 0.89 (0.40, 1.96) | 0.7698 |

To validate the results of the survival analyses, we looked at the performance of the new ICB against current standard of care predictive models by conducting logistic regression models where the binary outcome of biochemical recurrence or death was examined. We evaluated the Akaike information criteria (AIC) to conduct predictive model comparisons as a measure of the relative quality of the statistical models, as well as control for both goodness of fit and the complexity of the model. Given a set of candidate models for the data, the preferred model is the one with the minimum AIC value. The ICB has a significantly lower AIC than Gleason alone and RSG3 alone, both for BR (713.15 << 1016.69) and PCaSD (258.33 << 418.25) (Figure 4M). The same is true when incorporating the current clinico-pathologic parameters (seminal vesicle invasion, UICC staging, extracapsular extension, margins) into the model. The AIC for model with the ICB is much lower than Gleason with the current clinic pathologic parameters, both for BR (552.95 << 702.63) and PCaSD (233.7 << 308.17) (Figure 4N). The addition of the ICB significantly helps the model specification. This demonstrates that the interaction between Gleason and stromal grading provides more information than either marker alone in the model.

### Both stromal reactivity and tissue destruction are key factors in prostate cancer growth

To understand biological processes that could add more predictive information to our model, we examined tissue destruction *in silico*. This refers to the destruction of normal acinar structures present in the non-neoplastic prostate. As there are no techniques that can elucidate the specific nature of the initiating tumor cell in patients, we examined the correlation between the degree of tissue destruction by tumor cells (destruction of normal ducts and ECM) and the length of time a tumor takes to reach the edge of the simulation domain (years to maximal growth) (Figure 4O). The rate of tissue destruction is a composite function of basal and luminal epithelial cell death and loss of organization (ECM and BM reduction). We observed that 74% of the variation in the rate of growth could be explained by the degree of tissue destruction. The correlation between the rate of stromal activation and tumor growth (Figure 4P) improved this to 87%. The combination of both the degree of tissue destruction and the degree of stromal activation creates an almost perfect fit to predict the rate of tumor growth, with a correlation coefficient of 99% (Figure 4Q). These data suggest that the degree of tissue destruction could be used in conjunction with reactive stroma grading and/or the ICB, to predict PCa risk better, although currently there are no equivalent pathologic measurements that define tissue destruction. This will be an area of future development.

### Reactive stroma drives cancer evolution and heterogeneity

To better understand how the dialogue between tumor cells and stromal reactivity drives evolution in our mathematical representation, we examined two conditions of SR (Figure 5A-E). When a non-aggressive tumor cell (i.e. one with low rates of GF and MMP production) is seeded in our multiscale prostate peripheral zone model with a low SR, the rate of tumor evolution is faster than in a high SR environment (Figure 5F). Interestingly this does not translate into a larger tumor as the competition between the tissues (tumor and stroma) for the limited GF resource slows overall growth (Figures 5A-E). This competition, however, places a strong selection pressure upon the tumor, driving it towards cell phenotypes that produce more GF to survive. The increased selection pressure varies both spatially and temporally, depending on the relative local abundance of tumor and stromal cells. The resulting tumors tend be more phenotypically heterogeneous (Figure 5C). Conversely, the selection pressure for more aggressive phenotypes is much lower in tumor microenvironments characterized by high SR, even when the initial tumor cell is not aggressive (Figure 5E). The resulting tumors are less phenotypically heterogeneous and more driven by drift than selection. These tumors are the ones that grow and invade the fastest. This phenomenon is due to the abundance of stromal-derived GF not only within the tumor, but also further afield because of extended stromal activation. Of note is that the other evolving trait, MMP production, does not seem to be differentially selected between the high and low SR microenvironments (Figure S2). Although there is a trend for higher production in the high SR microenvironment, which may be due the more rapid growth these tumors display.

**Figure 5.**
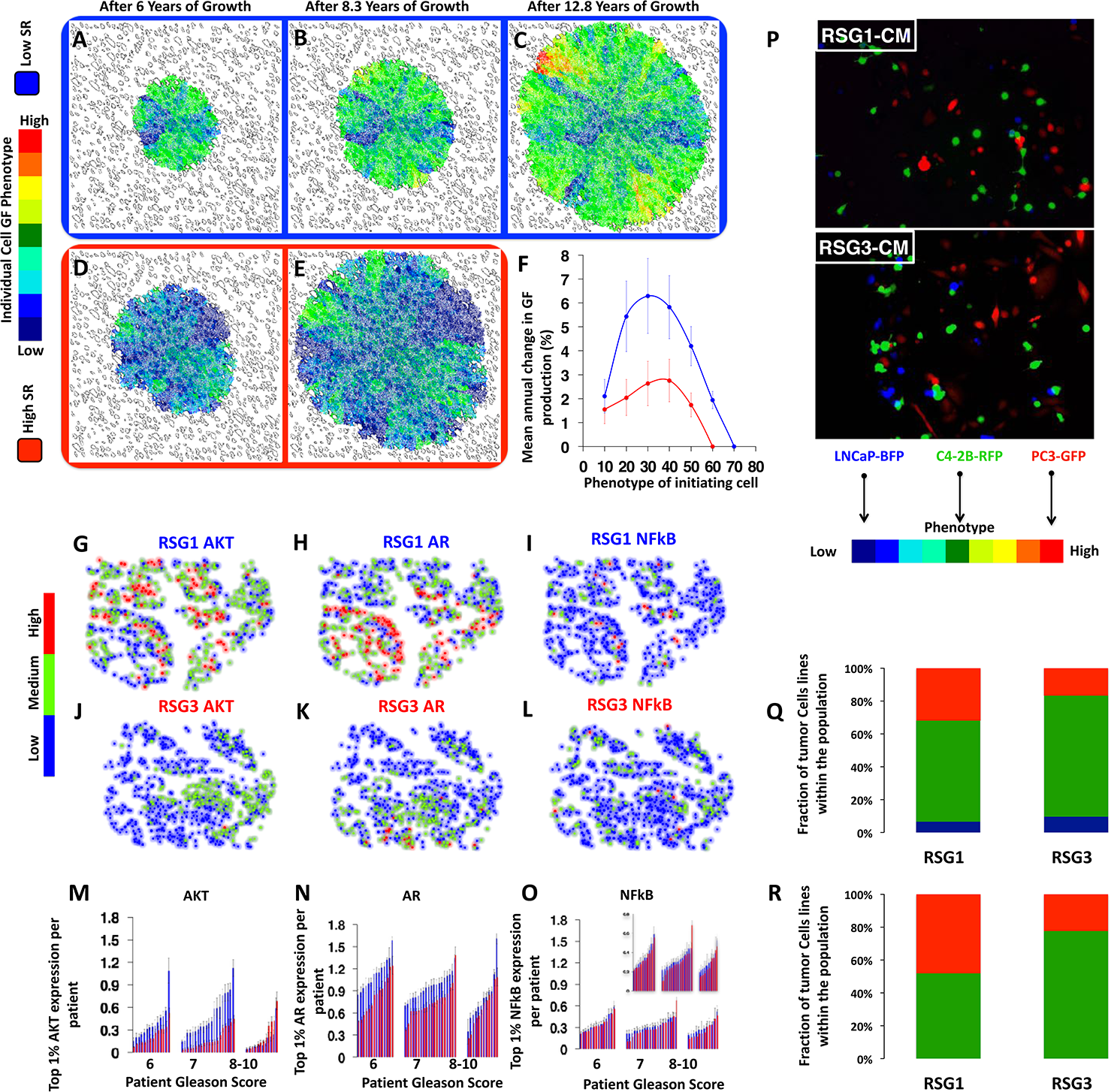
SR drives tumor cell evolution and progression: An in silico and clinical analysis. The evolution of tumor cell phenotypes through space and time (6-12.8 years) under low (blue) and high (red) SR conditions (A-E) (this figure extends the timescale of figure 2A-D). Heat map shows tumor cell phenotype (GF production) distribution in low SR (A-C) and high SR (D-E). Tumor cell phenotypic change in GF production from 8 different initiating phenotypes in high (red) and low (blue) SR environments (the average change and standard deviation across 300 simulations per initiating phenotype) is shown in panel F. Representative samples of triple immunostained biopsies for 3 well established PCa molecular regulators, phosphorylated AKT (G, J), Androgen receptor (AR) (H, K) and the central inflammatory regulator nuclear transcription factor kappaB (NFkB) (I, L) from two patients, one with RSG1 (blue border: G-I) and the other RSG3 (red border: J-L). Levels of expression were classified into low (blue), medium (green) or high (red). (M-O) Single cell quantitative analysis of the triple immunostained tissue sections for patients with RSG1 (blue) vs. RSG3 (red) in each Gleason category. The top 1% of gene expression in cells from each of the patient’s biopsies are shown (subset of the tumor cell population that would be under the greater selection pressure), each individual bar represents the average (and deviation) for a single patient over many cells. Prostate cancer cell lines LNCaP-BFP, C4-2B-RFP and PC3-GFP were cultured in the presence of conditioned medium (CM) from either NPF, RSG1-CAF or RSG3-CAF for 4 weeks. Representative images for each group are shown (P). Quantitation of individual cell populations was determined by FACS analysis (Q-R).

An intriguing prediction of our model is that the differences in tumor evolution are modulated in a non-linear way by the aggressiveness of the initiating phenotype (Figure 5F). In the midrange of initiating cell phenotypes (25-35), the difference in tumor evolution rates between high and low RS is greatest (Figure 5F), whereas the difference is minimized for the least and most aggressive initiating phenotypes (10 and 60 respectively). This is reminiscent of our previous findings where RSG3 has better predictive ability in tumors with a Gleason score of 7, compare to 8-10 (Yanagisawa et al., 2008).

To test this prediction in human samples, we performed an analysis of three well established prostate cancer molecular regulators: androgen receptor (AR), phosphorylated AKT1 and activated nuclear factor kappa B (NFκB) (phospho-p65). The analysis was performed on triple immunostained tissue sections of a large cohort of patients, previously described (Ayala et al., 2003), using image deconvolution, tissue compartment segmentation (i.e. tumor vs. stroma) and cell-by-cell analysis of each marker. Figure 5(G-L) shows tissue examples from patients with RSG1 (Figure G-I) vs. RSG3 (Figure K-L), where cell color reflects expression intensity for each of the individual cell expression of Phosphorylated AKT, Androgen receptor (AR) and the central immune regulator nuclear transcription factor B (NFκB).

Further analysis was performed on tissues from each Gleason category, and over RSG1 (Figure 5M-O, blue) and RSG3 (Figure 5M-O, red), like the ICB and *in silico* model predictions. We quantified AKT, AR and NFκB across all individual cells in tissues from a large population of patients (Figure 5M-O). Results of the average expression across all cells show a trend towards selection of a more aggressive phenotype in low RS (i.e. high levels of biomarker expression) (See supplemental Figure S3). To isolate a subset of the tumor cell population that would be under greater selection pressure, we analyzed the top 1% of expressing cells in each of the selected patients (Figure 5M-O). We identified a statistically significant difference between patients with RSG1 and RSG3, with higher expression of AR and AKT in RSG1, than in RSG3 in this cell population. Importantly, these differences were most apparent within the Gleason 6 and 7 patients, showing direct concordance with the *in silico* predictions (Figure 5F) where intermediate initiating cell phenotypes show the greatest evolutionary divergence.

We performed coculture experiments using three human prostate cell lines (LNCaP, C4-2B and PC3) representing mildly, moderately and aggressively invasive disease in medium conditioned by RSG1 or RSG3 tumor-derived fibroblasts (Figure 5 P-R). Equal numbers of cells either as pairs or all three lines together were plated and the number of each cell type at the end of the culture period was counted. These studies were consistent with the mathematical predictions in that the most aggressive cell type did not overgrow the other populations, and in most situations where all three lines were cocultured the dominant cell at the end of the study was the intermediate C4-2B line (Figure 5Q).

On inspection of some of the patient samples we noticed significant gradients in AKT and AR expression. We reasoned that such gradients maybe indicative of an evolving population. We developed a simple algorithm to examine the slope of expression across the triple stained samples by analyzing the change in expression spatially across each individual sample (Figure 6A), the larger the slope the more quickly expression changes with distance. We observe the largest slopes in Gleason 7 patients with RSG1 for all three markers (AKT, AR and NFκB) (Figure 6B). If we view the slope as a metric for evolution, this is again consistent with our in silico predictions, that the greatest evolution occurs at intermediate phenotypes with low SR (c.f. Figure 5F).

**Figure 6.**
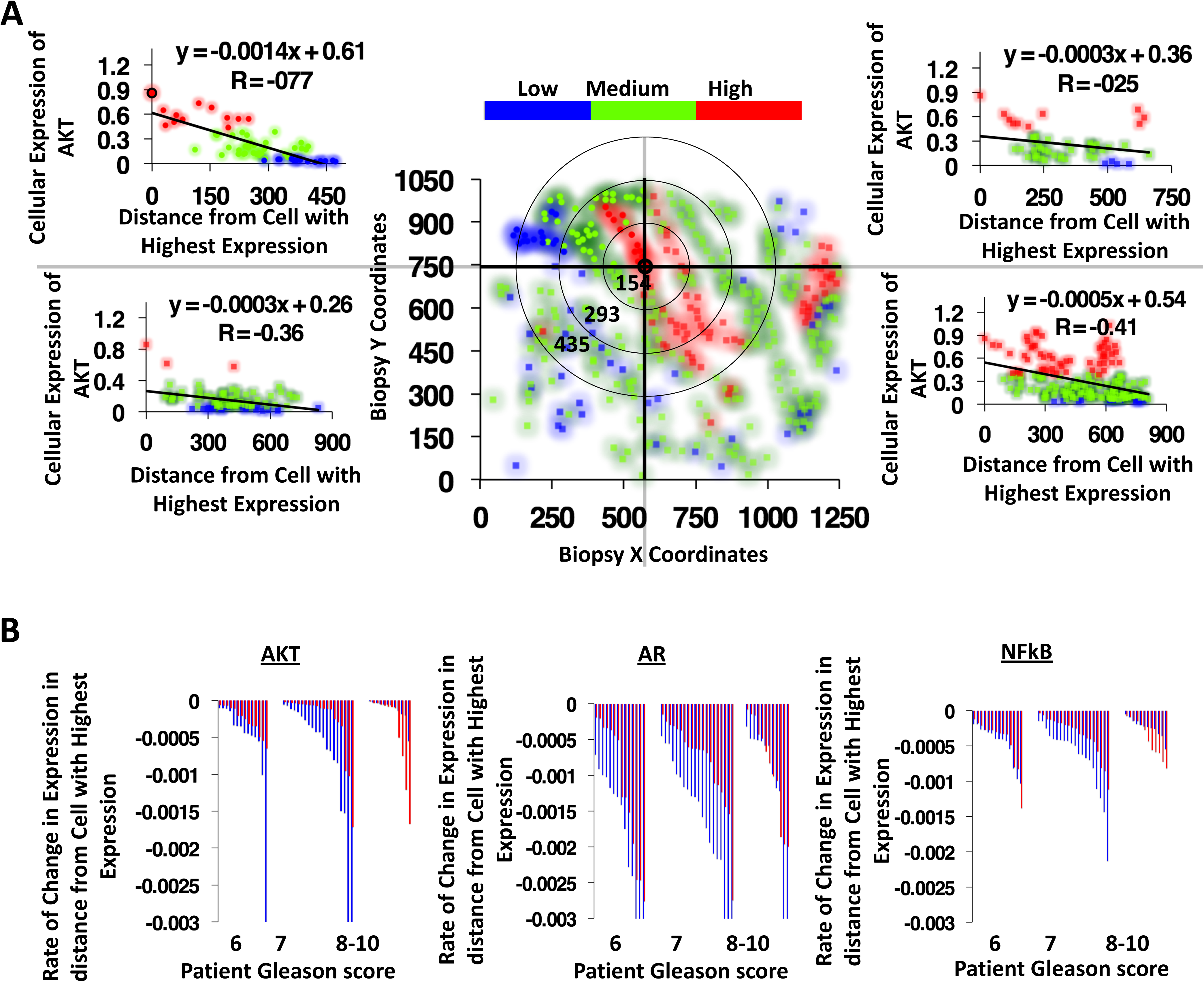
Calculating evolutionary gradients from patient biopsies. Analysis of triple stained tissue samples (Gleason 7 with RSG1) illustrating our approach to identify the most statistically significant evolutionary gradient in AKT expression. To identify the most significant gradient across a given biopsy, we analyzed the rate of change in expression through space starting from the cell with highest individual level of expression. Slope was calculated across radial distance from the cell with highest expression in the biopsy, in the example shown here, coordinates (575,746) (A). Analysis was performed on patients with RSG1 and compared to RSG3 in each Gleason category (B) showing the most significant slope per patient for the 3 molecular markers, AKT (left), AR (middle) and NF-B (phospho-p65, right). The larger the slope the more quickly expression changes with distance from the highest expressing cell.

## Discussion

*In vitro* and *in vivo* methods constitute the backbone of current biological knowledge. Yet they are limited in their ability to model complex interactions and diverse dynamics, due to their reductionist nature. Mathematical modeling readily captures the complex nature of cancer pathogenesis, both spatially and temporally. Critically, these models allow us to manipulate multiple factors both in isolation and simultaneously, combining key features as well as teasing apart their individual contributions, while still predicting how their interactions will shape tumor behavior. Features such as interactions at different scales and the stochastic nature of evolution can be tested experimentally *in silico*. Therefore, to better understand the dialogue between tumor and stroma we have implemented a hybrid multiscale model of carcinogenesis in the prostate peripheral zone. Results from this model generated novel concepts on the role of stroma in cancer evolution and led us to new ideas. Our model generated multiple hypotheses, some of which were tested *in vitro* and *in vivo* and subsequently validated in a large cohort of patients. This integrated paradigm merges data from biology and pathology on the microenvironmental regulation of PCa cell behavior and leads to novel insights in understanding cancer evolution, heterogeneity and progression. Our new *in silico* model is a significant step in the development of mathematical models that better mimic human prostate cancer.

Interactions between tumor stroma and epithelium are established regulators of prostate cancer progression (Dakhova et al., 2009; Hayashi and Cunha, 1991; Olumi et al., 1999). Cells derived from the reactive stroma (carcinoma associated fibroblasts – CAF) can drive tumor growth and invasion (Ao et al., 2007; Franco et al., 2011). In PCa the degree of stromogenic response to a tumor correlates to biochemical progression and cancer specific death (Ayala et al., 2003; Ayala et al., 2011). Our multiscale prostate peripheral zone model recapitulates these observations, demonstrating its ability to reproduce known biological and pathological processes and building confidence in its predictive power.

It is well established that stroma plays an important role in the development of prostate cancer. Our model goes beyond this finding and predicts that the interplay between the stroma and epithelium (i.e., the balance of individual cancer cell aggressiveness and stromal reactivity) is more important than either the stromal or epithelial properties alone. From our model, we have derived the following predictions: 1) the balance between stromal activation and tumor aggressiveness is key to tumor progression 2) stromal cells can enhance stromal activation, suggesting that reactive stroma can become self-activating under certain conditions, 3) tissue destruction is a potential biomarker of progression, and, 4) that the degree of stromal reactivity regulates tumor epithelial evolution. Our integrated approach allowed us to test and validate hypotheses 1, 2 and partially 4.

The model predicts that GFs produced by either the reactive stroma or cancer cells play complimentary roles and may compensate for each other. The significance of this balance was validated using both *in vivo* modeling and human studies. Results in both mice and humans were consistent with our mathematical prediction.

Concordantly, we identified that Gleason score and RSG quantitation (stromogenic carcinoma), when combined as the ICB, improves predictive power. In fact, the RSG quantitation significantly stratifies all the recurrence and death risk categories as suggested by the current Gleason Grading system. Due to its combinatorial nature, this system better addresses the issue of heterogeneity in both cancer differentiation and degree of stromal response. These results suggest that for patients whose tissue shows a high percentage of the tumor with stromogenic response, the risk should be considered higher than the standard Gleason assessment. Conversely, low percentage of RSG3 in a patient’s biopsy would imply a lower risk than the standard Gleason assessment. We show that the degree of stromal reactivity, when integrated with the current clinical in use methodology (Gleason with clinico-pathologic parameters), significantly improves PCa prognosis and may be useful to modify the way the patients will be treated. We would like to stress that the ICB can stratify the most of the problematic intermediate Gleason 7 category. It is also important to note that the percentage of tumor with RSG3 varies between Gleason subgroups. Gleason 6 requires a minimum of 21% of the tumor with RSG3 to be predictive. This increases to 61% and 71% in Gleason 7 and Gleason 8-10, demonstrating that the dynamic of the interplay between stroma and cancer has differential thresholds. This supports the balance concept between stroma and cancer cells identified in the mathematical model.

The most important prediction of this study is that evolutionary dynamics in tumors growing in the presence of stroma with high SR is different from those growing in association with low SR. The clear implication of this finding is that the progression of tumors reflects the ability of the host to respond to specific stimuli. In low SR situations, the rate of epithelial cell evolution is higher, leading to increased heterogeneity with selection for increased GF production. In contrast, high SR results in lower evolutionary pressures on the tumor epithelium, since sufficient GF is provided by the reactive stromal cells. Therefore, the differentiation state of the cancer cells tends to drift in terms of GF production, rather than undergoing rigorous selection for high expression needed for survival when an outside source is unavailable. This suggests that high levels of stromally-produced GF may result in less aggressive individual cancer cell phenotypes, but nonetheless the tumor constitutes a larger invasive mass. Using cell by cell data analysis of triple immunostained biomarkers in a cohort of human patients we could identify a signal that indicated that prostate cancer cells growing with low reactive stroma (RSG1) have a higher rate of selection, leading to evolution of a more advanced phenotype. These changes were more significant in Gleason 6 and 7 patients, than Gleason 8-10, (Figure 5G), concordant with our mathematical model predictions that identify greater evolution when starting with intermediate cancer cell phenotypes (Figure 5E). This makes biological sense, since more aggressive tumors (i.e. Gleason 8-10) may be less dependent on stroma for supplying GF, or they may require higher levels of stromal involvement to make a survival difference.

These evolutionary dynamics and associated tumor heterogeneity demonstrate why it is not possible to accurately assess patient risk by relying exclusively on tumor cell features (tissue architecture and molecular markers of individual cancer cells), especially for tumors with intermediate grades.

Our results therefore suggest two broad mechanisms that lead to invasive tumor growth. In the first case, the presence of highly reactive stroma within and surrounding the tumor provides excess GF, fueling growth. A positive feedback loop leads to a self-activating reactive stromal compartment, allowing tumor growth to continue indefinitely. This first case would correspond to high RSG and intermediate Gleason scores. In the second case, a lack of stromal reactivity instead promotes the evolution of tumor cell phenotypes favoring GF production. These altered cells, by nature, are less stromally-dependent and will eventually form invasive tumors. However, the tumor is relatively indolent until these GF-producing cells have developed and been selected, which takes time (perhaps even beyond the lifetime of the patient) when compared to the first case, where a tumor can immediately access high levels of GF.

In conclusion, our data indicate that the interactions between the tumor cells and reactive stroma shape the evolutionary dynamics of PCa and explain overall tumor aggressiveness (Figure 7). We show that the degree of stromal reactivity, when coupled with the current clinical biomarkers, significantly improves PCa prognosis, both for death and recurrence, that may alter treatment decisions. We also show that SR correlates directly with tumor growth but inversely modulates tumor evolution. This suggests that the aggressive stromal independent PCa may be an inevitable evolutionary result of poor stromal reactivity and that purely tumor centric metrics of aggressiveness may be misleading in terms on clinical outcome.

**Figure 7.**
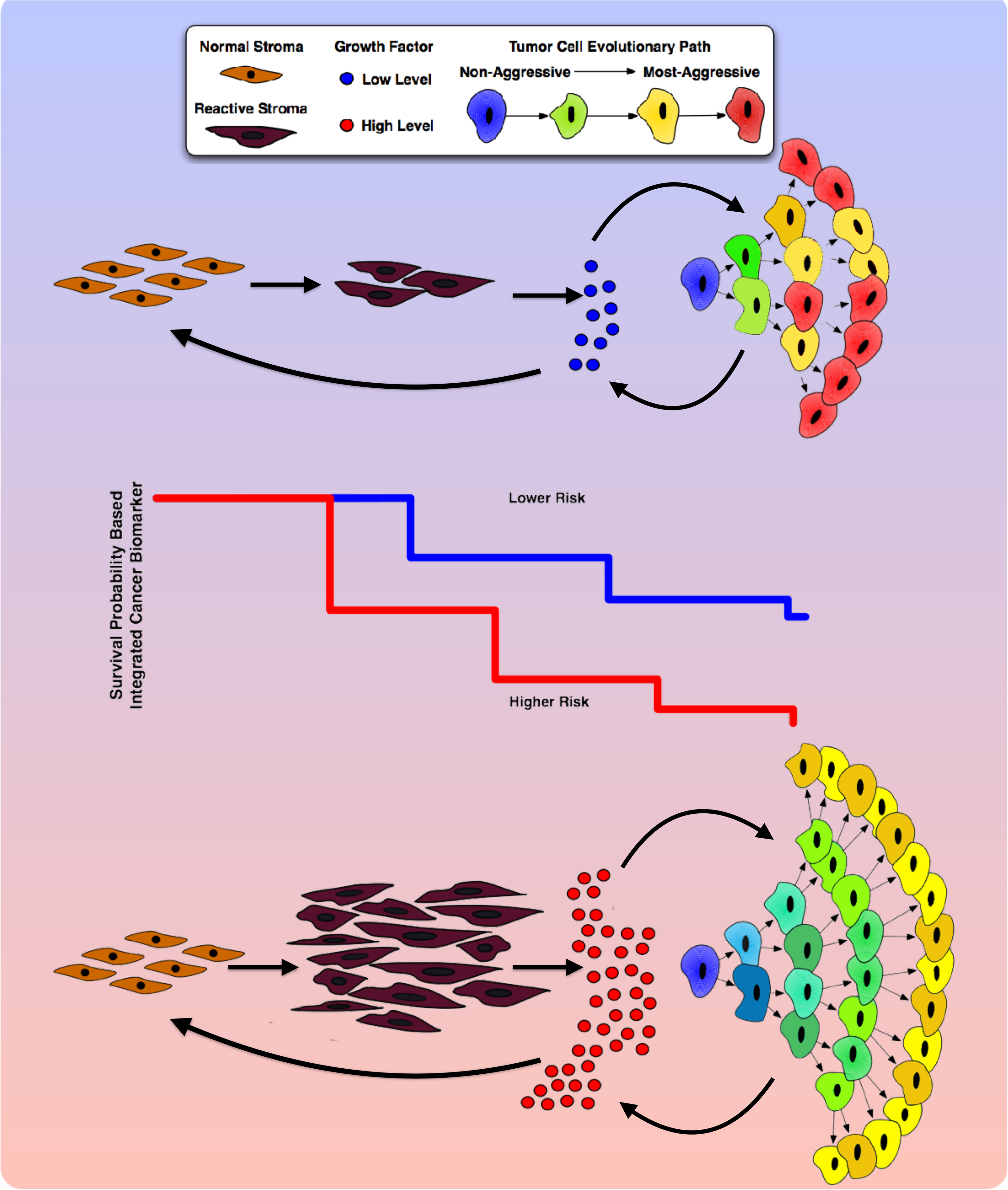
Interactions between tumor cells and stroma shape the evolutionary dynamics of PCa and drive overall tumor aggressiveness. Growth factor (GF) signaling is essential to both the tumor and the reactive stroma. Limited availability of GFs leads to an increased competition between tumor and stromal cells. This competition results in slower tumor growth (lower risk estimation) but also increased selection and more rapid evolution (leading to a more heterogeneous population). In contrast, where GFs availability is not limiting there are more mutualistic interactions between the tumor and stroma. This situation results in faster growing tumors (higher risk estimation) but, paradoxically, weaker selection pressure leading to less aggressive tumor cells (and a less heterogeneous population). Therefore, evolution of the most malignant phenotypes in a tumor cell population is not necessarily consistent tumor growth and invasion since it is modulated by the stromal response. In addition, tumor aggressiveness, as defined by Gleason score, is differentially modulated by stromal response. These different risk estimations and evolutionary dynamics (and tumor heterogeneity) mean that the overall behavior of patient tumors is driven by both the tumor cells (Gleason score) and the stromal response of the host (integrated cancer biomarker).

## Experimental Procedures

### Multiscale prostate peripheral zone model

The model we develop here is an extension of the hybrid cellular automaton (HCA) model described by Anderson and Basanta (Anderson, 2005; Basanta et al., 2009). The definition of an HCA model requires a set of partial differential equations (PDE), that characterize the physical microenvironment, a set life-cycle flowcharts that characterize the behavior of the cells under microenvironmental constrains (Figure 1E) and a cellular automaton framework to integrate them. The following system of non-linear PDEs define GF (*G*), MMP (*E*) and ECM/BM (*M*) as the three key continuous microenvironmental variables:

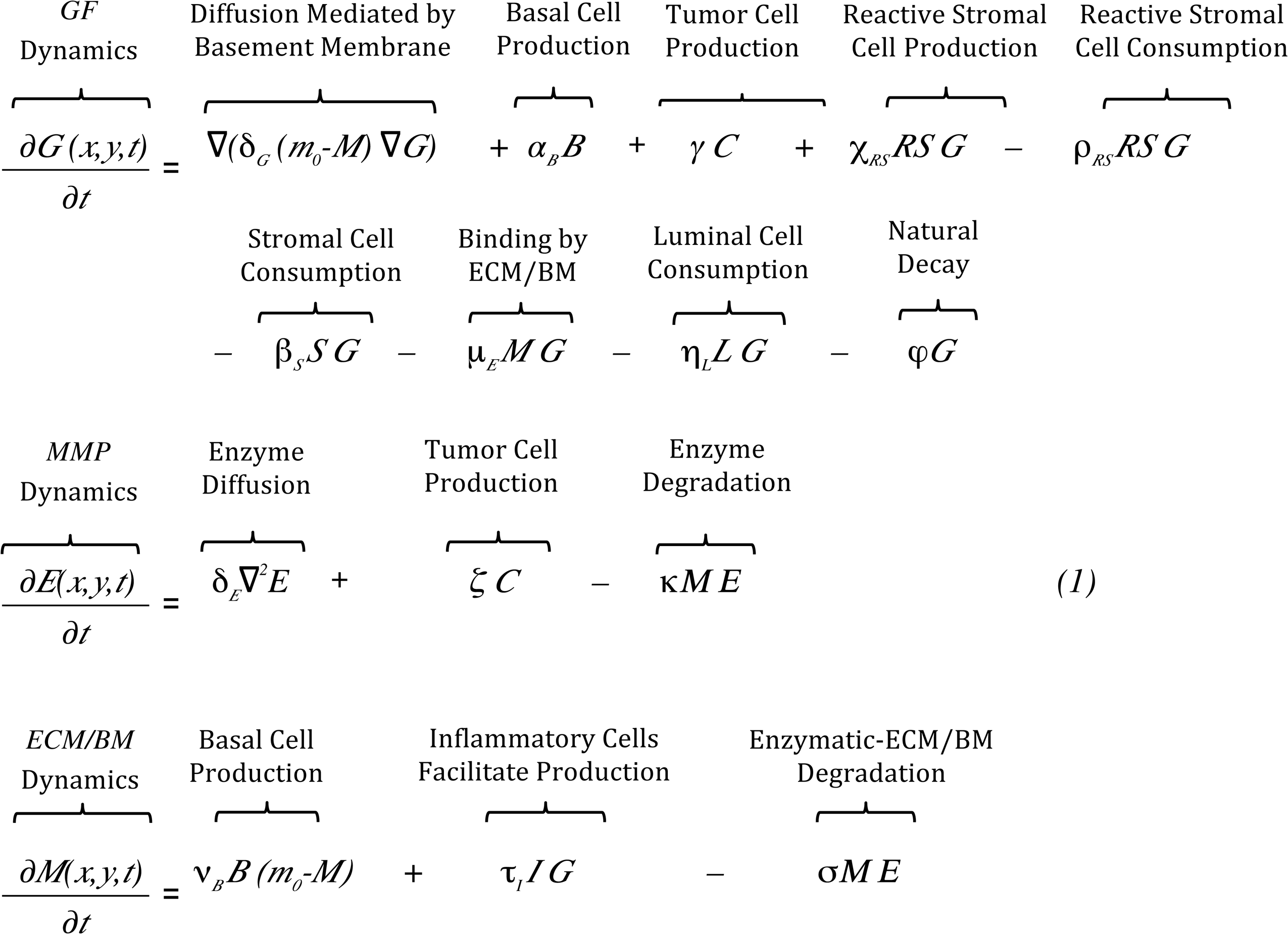

δ_*G*_ and δ_*E*_ are GF and MMP diffusion coefficients, respectively (Supplemental Information); *α_B_, γ,* χ_*RS*_, ρ_*RS*_, β_*S*_, μ_*E*_, η_*L*_, φ*, ζ,* κ, ν_*B*_, τ _*I*_, and σ are positive constants with biologically significant values as based on (Basanta et al., 2009). Then a discretized form of these equations is solved numerically on a 2-dimensional lattice that represents a small slice of prostate tissue. All cell types (tumor, basal, luminal, stromal and inflammatory) are modulated by the microenvironment on this lattice and can migrate, proliferate, die and mutate according to the life-cycle flowcharts (Figure 1E). Further detail can be found in Supplemental Information and Figure S1.

Tumor cells in the model have two continuously variable phenotypes, GF (*γ*) and MMP (*ζ*) production. These traits are passed from a parent tumor cell to its two daughter cells with some small variation, chosen at random from an interval equally weighted in both directions to avoid biased drift. The model is agnostic with respect to specific biologic mechanisms that underlie this drift, which could include gradual accumulation of mutations, regulation of gene transcription by epigenetics or aneuploidy, or changes in the number or structure of organelles, for example. The evolution and selection of these phenotypes in time and space is an important consideration of this work.

The switch between stroma (*S*) and reactive stroma (*RS*) phenotypes is driven by GF stimulus. Reactive stromal cells are activated if the level of GF (*G*) is above the threshold *G*_RS_, and are deactivated if the growth factor level *G* falls below the threshold *G*_RS_.

### Clinical Specimens

To validate the findings of the mathematical model and its biologic validation, we used a unique human tissue resource. The cohort was selected from patients operated by a single surgeon, without additional forms of therapy other than surgery, and over 20 years of follow-up.

### Cohort Enrollment and Follow-up

The Baylor Prostate Cancer database contains information on over nine thousand patients who underwent radical prostatectomies at one of the Baylor College of Medicine affiliated institutions and provided tissues (IRB H-1158). Of these patients, 1,291 were operated on by a single surgeon, between 1983 and 1998 without any previous form of adjuvant therapy such as radiation or hormonal therapy. This study was approved by the Baylor Institutional Review Board (IRB H-11436).

Radical prostatectomy specimens from these patients were processed using whole-mount slides according to procedures described previously (Wheeler and Lebovitz, 1994). After surgery, the prostate specimens were sliced into 5 mm-thick tissue whole mounts. The tissue slices were then fixed in 10% neutral-buffered formalin and embedded in paraffin according to a routine procedure. A single pathologist performed the pathological analysis that included staging, pathological stage, margins, capsular penetration, seminal vesicle invasion, primary and secondary Gleason grades from biopsy and prostatectomy, lymph node status, tumor volume, and geographic location. The clinical and pathological data of patients who met the entry criteria were available for analysis. The clinical follow-up data include prostatic-specific antigen recurrence (defined as prostatic-specific antigen 0.4ng or two consecutive rises), clinical metastasis, and prostate cancer specific death.

#### Clinical Characteristics

Age of patients ranged from 37 to 80 with a mean of 62 and median of 63 years. The patients were followed postoperatively for an average of 42.08 33.2 months (mean SD, median 45.2, maximum 167.74). Preoperative prostatic-specific antigen level was available in 603 prostate cancer cases and ranged from 0.3 to 100 ng/mL with a median of 7.2 ng/mL, and a SD of 10.99 ng/mL. Approximately 30% of the patients had a preoperative-prostatic-specific antigen level 10.5 ng/mL. Approximately 7% had a Gleason score 6; 85% had Gleason score of 6 or 7, and 8% had a higher Gleason score (8 to 10). Lymph node metastasis was found in 40 (6.4%) patients, and biochemical recurrence was seen in 120 patients (19.3%). Extra-capsular extension was found in 44.5%; margins were positive in 15.3%, and seminal vesicle invasion had occurred in 12.4% of the patients.

Perineural Invasion Diameter was obtained, as described in Maru et al. (Maru et al., 2001). Quantification of stromogenic carcinoma pattern (RSG) was performed as previously described (Ayala et al., 2011).

### Statistical Analysis

Cox proportional hazard regression analyses for each of biochemical recurrence and PCa specific death were conducted to evaluate univariable and multivariable association between a stromal marker (i.e., the percentage of RSG3 within radical prostatectomy PCa) and risk of recurrence or death. Important standard clinical-pathological risk factors such as Gleason grade, extra-capsular extension, seminal vesicle invasion, margins, lymph node status and preoperative PSA were also considered while developing multivariable models to examine the contribution of a stromal marker (i.e., %RSG3) over traditional factors. Specifically, we focused on determining the effect of stromal marker addition on epithelial cancer marker (i.e., Gleason grade) by evaluating whether the association between Gleason score and risk of recurrence (or death) differs by levels of % RSG3, which led to a combining of these two markers resulting in the integrated cancer biomarker (ICB). While developing integrated marker, assessment of linearity/functional form and residuals (Therneau and Grambsch, 2000) was made to ensure underlying linearity assumptions between the predictors and outcome are valid. To establish optimal cut-off values of %RSG3 for every Gleason category, we used minimum P value approach as well as goodness of fit tests for overall significance of the model (Grønnesby and Borgan, 1996). A test of proportional hazards assumption was also performed and indicated that there was no statistically significant evidence of assumption violation (Grambsch et al., 1995; Therneau and Grambsch, 2000).

To validate our findings from the survival analyses, we looked at the predictive performance of the new ICB against current standard clinical-pathological risk factors by conducting multivariable logistic regression models, where the binary outcome of biochemical recurrence or death was examined. The prediction accuracy of each model was measured using area under the receiver operating characteristic (ROC) curve (AUC). Given the fact that increase of AUC was very small when model already includes the highly significant standard risk factors (e.g., AUC >0.85), the magnitude of improvement in AUC may not be nearly as meaningful as the value of AUC itself (Pepe et al., 2004). Therefore, in addition to AUC, we considered deviance-based measures such as Akaike information criteria (AIC), which is widely used to examine whether the addition of an extra independent variable improves the model specification (i.e., the relative quality of model), while controlling for goodness of fit and the complexity of the model at the same time. Hosmer-Lemeshow goodness-of-fit test was also constructed to examine agreement between observed outcomes and predictions. All Analyses were performed primarily using widely available tools in SAS^®^ version 9.4 (SAS Institute, Cary, NC) at a significance level of 0.05.

### AR, pAkt and pNF-κB p65 (phospho S276) triple staining

New TMA slides were triple immunohistochemicaly stained by using the AR (Biocare Medical, cat# CM109A), pAkt (Dako, cat# M3628) and pNF-κB p65 (phospho S276) (Abcam, cat# ab106129) antibodies. Before the triple staining, all 3 antibodies were tested in test tissue microarray slides containing different human prostate tissue samples. Briefly, sections were deparaffinized in xylene, rehydrated through decreasing concentrations of alcohol ending in PBS, subjected to heat-induced antigen retrieval in Dako’s Target Retrieval Solution (pH 9.0, cat# S2367) for 4 minutes, 125°C in a Pascal instrument (Dako cat# S280030), and allowed to cool off at room temperature. Endogenous peroxidase activity was quenched in 3% hydrogen peroxide solution in distilled water for 10 minutes at room temperature. To inhibit non-specific staining, sections were incubated with a protein blocking solution (Dako cat# X0909) for 10 minutes at room temperature, then incubated with the mixture of mouse monoclonal antibody against AR (1:100) and rabbit polyclonal antibody pAkt (1:10) in antibody diluent (Dako, cat# S0809), 1 h at room temperature. Sections were washed and the bound antibodies were detected by using the Biocare Medical MACH2 Double Stain 2 Polymer Detection kit (mouse-HRP + rabbit-AP, cat# MRCT525), with diaminobenzidine (DAB, for AR) and Vulcan Fast Red (for Akt, Biocare Medical cat# FR805) as chromogens.

To ensure that the third antibody staining will not cross react with the AR and pAkt staining, sections were incubated with a denaturing solution for 2 minutes (Biocare medical Cat# DNS001), then incubated with rabbit polyclonal antibody against pNF-κB p65 (phospho S276) (1:300), 2 h at room temperature. Sections were washed and the bound antibody was detected by using a Biocare Medical MACH 2 Rabbit HRP-Polymer (Cat# RHRP520) with Vina Green (Biocare Medical, cat# BRR807AH) as chromogen.

Finally, sections were counterstained with Cat hematoxylin (1: 4 diluted with water, 40 seconds, Biocare medical Cat# CATHE) and the bluing reagent (10 seconds, Statlab Medical Products cat# SL203), then air dried at 65°C for 15 minutes, mounted with EcoMount (Biocare medical cat# EM897L).

### Quantification of AR, pAkt and pNF-κB p65 using image deconvolution, segmentation and analysis

Hot spot areas of expression of the 3 biomarkers within the PCa tissues were imaged using the Vectra 1.4.0 (one 200x images per core). A combination of deconvolution imaging (such as Vectra®) and image segmentation technology (such as INFORM® 2.1.1) was utilized.

All immuno stained slides were digitized with the use of a multispectral imaging system which enabled capturing a series of images from a single field at spectrum of specific wavelengths (420nm to 720 nm). Multiple series of images taken at different wavelength at one shut is called “image cube”. Image cubes were created for every case and saved in both multispectral .im3 and JPEG formats. All images were taken at 200x magnification, and capture more than 95% of 0.6 mm tissue cores. The measurement of image spectral wavelengths enables more accurate separation of the tissue, and cellular components.

The InForm image segmentation system was used to separate non-neoplastic and neoplastic PCa tissues from the normal muscular host stromal tissues, as well as reactive stroma. Signal was analyzed only in the epithelial component.

Image segmentation software was used for tissue and cellular analysis of the tumoral stroma in the prostatic adenocarcinoma. Pictures from each case were reviewed individually and only tumor and tumoral stroma areas were selected for further analysis.

Tissues were algorithm segmented into compartments (cancer epithelium and cancer stroma); each compartment segmented into individual cells and each cell segmented into nuclei and cytoplasm. Individual cells were recognized within each compartment and AR, pAkt and pNF-κB p65 signal was separated and analyzed in each compartment of the tumor separately in each individual cells. All were analyzed in the cytoplasm of the cancer cells. In conclusion, we were able to provide readouts per individual cells, in each compartment, each with coordinates for spatial localization within the tissues.

### Isolation and Characterization of Carcinoma Associated Fibroblasts

Isolation and validation of the tumor-inducing abilities of the CAF from tissue samples was performed as previously described (Olumi et al., 1999). Stromogenic classification of CAF isolated from patients was performed by scoring of Masson’s trichrome stained tissues according to standard guidelines (Ayala et al., 2003). Briefly, tumors with reactive stroma comprising 5% to 15% of the tumor were classified as RSG1, 15% to 50% RSG2, whereas those with more than 50% RSG3. Concurrently samples were scored for Gleason grade. Immunohistochemical staining for Ki67 was performed and a labeling index calculated for each sample.

### Effects of RSG status of source material on the tumor-inductive capability of CAF cells

To test the effects of RSG characteristics on tumor growth, an initiated but non-tumorigenic prostate epithelial cell line BPH1 was recombined with CAF with different RSG and grafted under the renal capsule of castrated, testosterone supplemented CB17Icr/Hsd-severe combined immunodeficient (SCID) mice (Harlan) as previously described (Ao et al., 2007). The BPH1 cell line responds to the pro-tumorigenic effects of CAF by undergoing malignant transformation (Olumi et al., 1999). We had previously noted that the volume of such tumors varies by the patient source of the CAF but had not examined this phenomenon formally. CAF from each RSG group were used in a standardized recombination assay and grafted to SCID mice for 12 weeks. The resultant tumors were harvested, and their area in a central cross section and invasiveness measured from the surface of the kidney to the deepest point of penetration was determined.

To test how PCa cell lines respond to CAF, and specifically to determine whether such a response is a function of the source tumor from which the CAF were derived, three cell lines (LNCaP, C4-2b and PC3) representing progressively increasingly aggressive disease were combined with CAF from various RSG-defined tumors and tested in tissue recombination experiments in SCID mice, as described above.

### Testing Stromal Auto Activation

Conditioned media using RPMI 1640 supplemented with 0.5% fetal bovine serum and 10^-8^ M testosterone was collected after 48-72 hours from: two sets of cancer-associated fibroblasts and control BHPrS1-EV cells.

BHPrS-EV (∼p5) cells were plated in a six-well dish at a starting density of 50,000 cells/well. After 24 hours, to allow attachment, the growth medium was changed to one of the experimental conditioned media. Media were changed twice a week and cells were subcultured as needed. Cells were harvested, RNA was isolated (BioRad RNA Isolation kit and cDNA kit), and qPCR was performed at the end of each week to assess various markers of activated stroma including CD90, TGFß1, SDF1, and αSMA.

### Effects of stromal conditioned medium on the growth

Normal prostate fibroblasts (NPF) and cancer-associated fibroblasts derived from RSG1, RSG3 tumors were grown in RPMI 1640 supplemented with 0.5% fetal bovine serum and 10^-8^ M testosterone for 48-72 hours, after which the media were collected, centrifuged, and filtered (0.45μm).

PC3, C4-2B and LNCaP cells (from ATCC and MD Anderson) representing progressively less aggressive tumors, were transduced with lentiviral constructs carrying fluorescent color tags (colors were a gift from Dr. Andreis Zjilstra, Vanderbilt University Medical Center). Cells were grown under positive Blastocydin selection and then moderate color expression was selected using FACS, to generate green, red and blue lines designated: PC3-GFP, LNCaP-BFP, and C42B-RFP. Cells were seeded (all combinations mixtures of single cell line alone; the three pair combinations and with all three colors together) in 96-well dishes in triplicate at an initial density of 5,000 total cells per well. The cells were allowed 24 hours to attach after which the medium was changed to a 1:1 mixture of the CAF-CM and RPMI 1640, serum adjusted to 0.5% FBS. Medium was changed twice weekly for 4 weeks and images were taken at the end of each week to determine the distribution of cells. At the termination of the experiment the cells were passed through a MACS Quant FACS analyzer (Miltenyi) to generate color-specific counts of total cells per well.

